# An ALS-associated KIF5A mutant forms oligomers and aggregates and induces neuronal toxicity

**DOI:** 10.1101/2022.03.28.486133

**Authors:** Juri Nakano, Kyoko Chiba, Shinsuke Niwa

## Abstract

KIF5A is a kinesin superfamily motor protein that transports various cargos in neurons. Mutations in *Kif5a* cause familial amyotrophic lateral sclerosis (ALS). These ALS mutations are in the intron of *Kif5a* and induce mis-splicing of KIF5A mRNA, leading to splicing out of exon 27, which in human KIF5A encodes the cargo-binding tail domain of KIF5A. Therefore, it has been suggested that ALS is caused by loss of function of KIF5A. However, the precise mechanisms regarding how mutations in KIF5A cause ALS remain unclear. Here, we show that an ALS-associated mutant of KIF5A, KIF5A(Δexon27), is predisposed to form oligomers and aggregates in cultured mouse cell lines. Interestingly, purified KIF5A(Δexon27) oligomers showed more active movement on microtubules than wild type KIF5A *in vitro*. Purified KIF5A(Δexon27) was prone to form aggregates *in vitro*. Moreover, KIF5A(Δexon27)-expressing *Caenorhabditis elegans* neurons showed morphological defects. These data collectively suggest that ALS-associated mutations of KIF5A are toxic gain-of-function mutations rather than simple loss-of-function mutations.

## Introduction

Neuronal development, function, and maintenance depend on intracellular transport (Hirokawa, Noda, Tanaka, & Niwa, 2009). Kinesin superfamily proteins (KIFs) and cytoplasmic dyneins are molecular motors that enable anterograde and retrograde transport, respectively (Hirokawa, Niwa, & Tanaka, 2010; Kardon & Vale, 2009). Among KIFs, the Kinesin-1, -2, and -3 family members are the main anterograde transporters in neurons (Hall & Hedgecock, 1991; Niwa, Tanaka, & Hirokawa, 2008 ; Okada, Yamazaki, Sekine-Aizawa, & Hirokawa, 1995; Scholey, 2008; Vale, Reese, & Sheetz, 1985; Xia et al., 2003). KIFs generally consist of a conserved motor domain, a dimerized coiled-coil domain, and a cargo-binding tail domain. The motor domain exhibits microtubule-stimulated ATPase activity, which allows the protein to move along microtubules (Hackney, 1995). Each KIF has a specialized tail domain (Vale, 2003) that binds to specific cargo vesicles and protein complexes (Hirokawa et al., 2009).

Advances in genomic sequencing technology have allowed identification of many disease-associated mutations in motor protein genes (Hirokawa et al., 2010; Holzbaur & Scherer, 2011). Mutations in KIFs and dynein subunits often cause motor neuron diseases. For example, mutations in Kinesin-3 family members, such as KIF1A and KIF1B, cause hereditary spastic paraplegia and Charcot-Marie-Tooth disease type 2A1 (Boyle et al., 2021; Budaitis et al., 2021; Chiba et al., 2019 ; Zhao et al., 2001). Mutations in dynein heavy chain 1 (DYNC1H1) cause Charcot-Marie-Tooth disease type 2O and spinal muscular atrophy-1 (Harms et al., 2012; Weedon et al., 2011). Mutations in the p150 subunit of dynactin (DCTN1), an activator of dynein, cause amyotrophic lateral sclerosis (ALS) (Munch et al., 2004). Gain-of-function mutations in BICD2, a cargo adaptor protein for the cytoplasmic dynein complex, also cause spinal muscular atrophy (Huynh & Vale, 2017). KIF5A mutations are associated with hereditary spastic paraplegia (SPG) and ALS (D. Brenner et al., 2018; Nicolas et al., 2018; Reid et al., 2002). KIF5A transport protein complexes and membrane organelles such as neurofilaments, RNA granules and mitochondria (Hirokawa et al., 2010; Kanai, Dohmae, & Hirokawa, 2004; Uchida, Alami, & Brown, 2009 ; Xia et al., 2003). The mutated residues differ between KIF5A-associated SPG and ALS. SPG is caused by motor domain mutations in KIF5A that abolish the motor activity of the KIF5A motor (Ebbing et al., 2008). However, ALS-associated KIF5A mutations commonly induce mis-splicing and deletion of exon 27 (Fig 1A) (D. Brenner et al., 2018; Nicolas et al., 2018). The Δexon27 mutation induces frameshift, leading to abnormal C-terminal tail (Fig 1A). Previous studies have suggested that ALS-associated mutations in KIF5A are loss-of-function mutations because of the deletion of the cargo-binding tail domain (D. Brenner et al., 2018; Nicolas et al., 2018). However, the precise molecular mechanism of KIF5A-associated ALS has not been shown.

**Figure 1.**
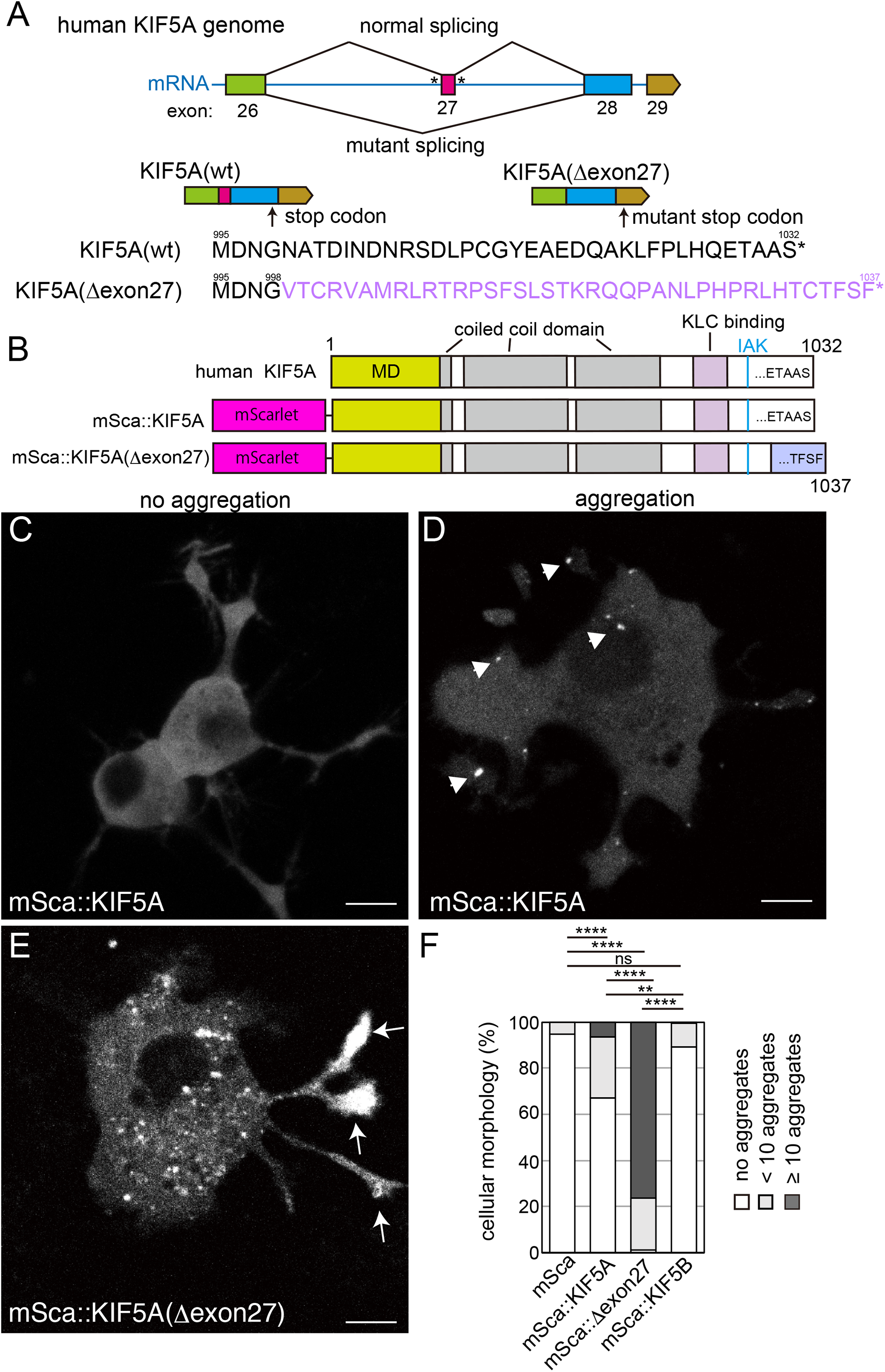
Cellular distribution of amyotrophic lateral sclerosis (ALS)-associated KIF5A mutations. (A) Schematic drawing of ALS mutations in KIF5A. ALS-associated KIF5A mutations (asterisks) are often found in introns around exon 27 and commonly induce mis-splicing of exon 27 and frameshifts. Amino acid sequences show the C-terminal sequences of wild type KIF5A (black font) and KIF5A(Δexon27) (purple font). (B) Schematic drawing showing mScarlet-fused wild type KIF5A (mSca::KIF5A) and mScarlet-fused mutant KIF5A(mSca::KIF5A(Δexon27)). Motor domain (MD) (1-335a.a.), coild-coil domains (331 - 374 a.a., 408 - 539 a.a. and 632 - 800 a.a.), light chain binding domain (KLC binding) (822 - 905 a.a) and IAK motif (918-920 a.a.), that is required for the autoinhibition, are shown. (C–F) mSca::KIF5A and mSca::KIF5A(Δexon27) were expressed in CAD cells and analyzed by fluorescence microscopy. (C and D) Representative images showing the localization of mSca::KIF5A (C and D) and mSca::KIF5A KIF5A(Δexon27) (E). Arrow heads and arrows indicate cytoplasmic aggregates and aggregates accumulated to neurite tips, respectively. Bars, 10 μm. (F) The cells were classified by the mScarlet signal distribution into three categories: no aggregates (white), no more than 10 aggregates (light gray) and more than 10 aggregates (dark gray), and the number of cells in each category was counted. N = 98 mScarlet-expressing cells, 112 KIF5A(wt)-expressing cells, 101 KIF5A(Δexon27)-expressing cells, and 177 KIF5B-expressing cells. A chi-square test with Bonferroni correction was used for data analysis. ns, adjusted P value > 0.05 and not statistically significant. *, adjusted P value < 0.05. **, adjusted P value < 0.01, **** adjusted P value < 0.0001.

We show here that the product of ALS-associated KIF5A alleles, KIF5A(Δexon27), is predisposed to form oligomers and aggregates. Binding of KLC1, a cargo adaptor of KIF5A, was not affected by Δexon27. Interestingly, KIF5A(Δexon27) oligomers showed higher motility than those of wild type KIF5A regardless of the presence of KLC1. Furthermore, exogenous expression of KIF5A(Δexon27) caused defects in the neuronal morphology of *Caenorhabditis elegans*. Collectively, these findings suggest that ALS-associated mutations in KIF5A are toxic gain-of-function mutations rather than simple loss-of-function mutations.

## Results

### KIF5A(Δexon27) forms aggregates in the cell

To study the molecular mechanism of ALS caused by KIF5A mutations, we expressed mScarlet-fused KIF5A (mSca::KIF5A) and mScarlet-fused KIF5A(Δexon27) (mSca::KIF5A(Δexon27)) in a neuron-like cell line, CAD (Qi, Wang, McMillian, & Chikaraishi, 1997) (Fig 1B–F). 20 hours after the transfection, mSca::KIF5A was mostly diffuse throughout the cell, but approximately 30% of mSca::KIF5A-expressing cells showed small aggregates in the cytoplasm (Fig 1C, D, and F), which is consistent with our prior finding showing a propensity of KIF5A to form oligomers (Chiba, Ori-McKenney, Niwa, & McKenney, 2022). The proportion of cells showing aggregation was increased among cells expressing mScarlet-fused KIF5B, a homologue of KIF5A (Hirokawa et al., 2009). Only 10% of mSca::KIF5B-expressing cells exhibited aggregates. In contrast, mSca::KIF5A(Δexon27) formed many aggregates in the cytoplasm, as noted in 97% of cells. We observed that aggregates often accumulated in the tip of neurites in CAD cells (Fig 1E arrows), which is similar to the localization of KIF5A with constitutive-active mutations (Nakata, Niwa, Okada, Perez, & Hirokawa, 2011). Few aggregates were found in mScarlet-expressing cells (Fig 1F). Formation of aggregates is not due to the higher expression of mSca::KIF5A(Δexon27) because most mSca::KIF5A(Δexon27)-expressing cells had aggregates even 6 hours after the transfection. These data suggest that KIF5A is predisposed to form aggregates in the cell and that the ALS-associated mutation Δexon27 strongly enhances aggregate formation.

### KIF5A(Δexon27) co-aggregates with wild type motors

We investigated whether the ALS-associated KIF5A protein co-aggregates with wild type motors. mSca::KIF5A(Δexon27) was co-expressed with enhanced green fluorescent protein (EGFP)-labelled KIF5A (EGFP::KIF5A). Previous studies have shown that KIF5A forms a homodimer and moves along microtubules (Hackney, 1995; Hackney, Levitt, & Wagner, 1991; Vale et al., 1996). As expected, mSca::KIF5A(Δexon27) co-aggregated with EGFP::KIF5A(wt) in the cell (Fig 2A–D), and aggregates accumulated at the tips of CAD cell neurites. *Homo sapiens* have three homologous kinesin heavy chain genes: KIF5A, KIF5B, and KIF5C (Hirokawa et al., 2009). These three proteins are highly conserved but do not form heterodimers (Kanai et al., 2000). Interestingly, mSca::KIF5A(Δexon27) co-aggregated with EGFP::KIF5B(wt) (Fig. 2E–H). However, EGFP::KIF5B was diffuse in the cytoplasm when mSca::KIF5A did not form aggregates (Fig 2E and F), and EGFP::KIF5B(wt) uncharacteristically accumulated in neurite tips and co-aggregated with mSca::KIF5A(Δexon27) (Fig 2G and H). In contrast, mSca::KIF5A(Δexon27) did not co-aggregate with EGFP alone (Fig 2I and J). These data suggest that KIF5A(Δexon27) co-aggregates with wild type KIF5A and KIF5B motors in the cell.

**Figure 2.**
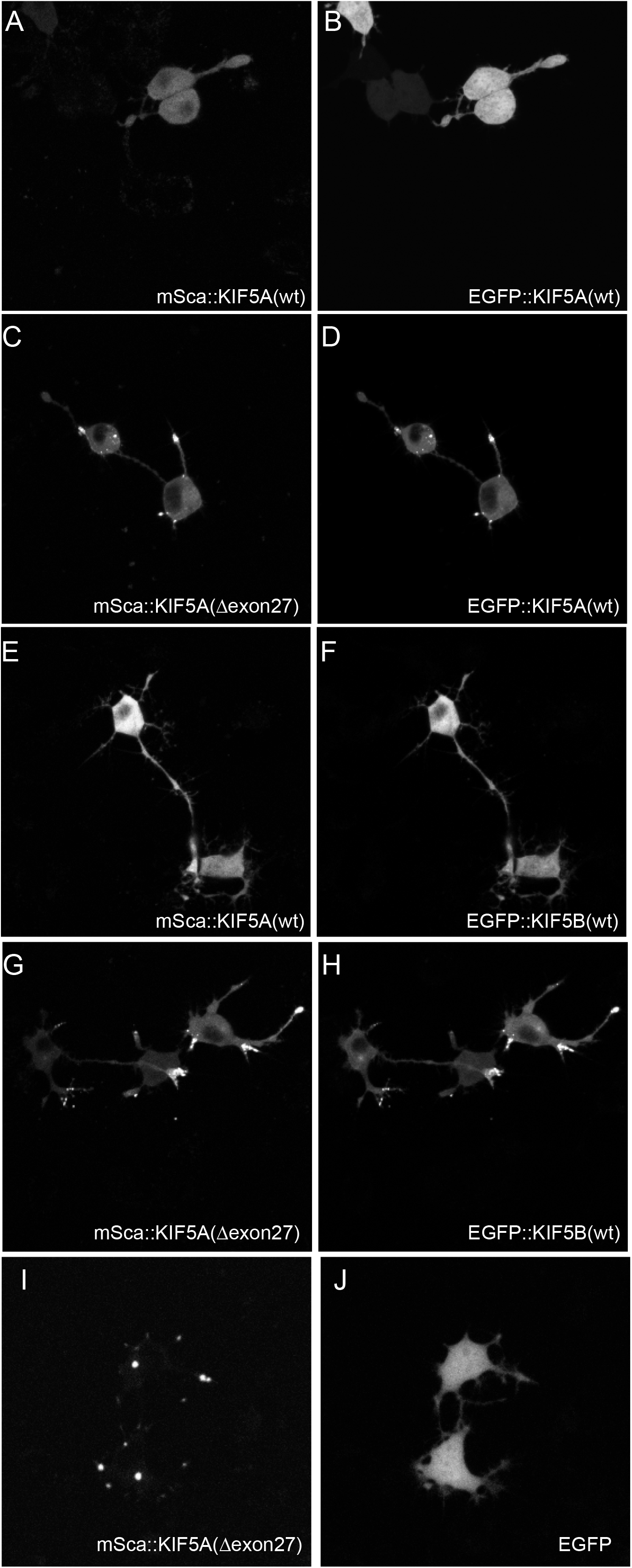
Co-aggregation of amyotrophic lateral sclerosis-associated KIF5A. mScarlet fused to KIF5A or KIF5A(Δexon27) was co-expressed with enhanced green fluorescent protein (EGFP)-fused proteins in CAD cells. (A and B) Representative images showing mSca::KIF5A (A) and EGFP::KIF5A (B) co-expressing cells. When mSca::KIF5A did not form aggregates, EGFP::KIF5B did not co-aggregate in the same cell. No cells (0/70) showed co-aggregation. (C and D) Representative images showing mSca::KIF5A(Δexon27) (C) and EGFP::KIF5A (D) co-expressing cells. Note that EGFP::KIF5A co-aggregated with mSca::KIF5A(Δexon27). All cells (85/85, 100%) showed co-aggregation. (E and F) Representative images showing mSca::KIF5A (E) and EGFP::KIF5B (F) co-expressing cells. When mSca::KIF5A did not form aggregates, EGFP::KIF5B did not co-aggregate as well. No cells (0/70) showed co-aggregation. (G and H) Representative images showing mSca::KIF5A(Δexon27) and EGFP::KIF5B co-expressing cells. Note that EGFP::KIF5B co-aggregated with mSca::KIF5A(Δexon27). All cells (82/82, 100%) showed co-aggregation. (I and J) Representative images showing mSca::KIF5A(Δexon27) and EGFP co-expressing cells. Even when mSca::KIF5A(Δexon27) formed aggregates in the cytoplasm, no GFP aggregation was observed. No cells (0/54) showed co-aggregation. Bars, 10 μm.

### KIF5A(Δexon27) oligomerizes *in vitro*

Kinesin-1 is a heterotetramer composed of two heavy chains (KIF5) and two light chains (KLC) (Hackney et al., 1991). We have previously shown that KIF5A has a propensity to form oligomers *in vitro* (Chiba et al., 2022). To examine the effect of the Δexon27 mutation on the interaction of KIF5A with KLC1 and oligomerization, we next purified heavy chain dimers (KIF5A) and Kinesin-1 tetramers (KIF5A-KLC1) with or without the Δexon27 mutation. First, we expressed full-length KIF5A::mSca and KIF5A(Δexon27)::mSca in sf9 cells and purified them by affinity chromatography and size exclusion chromatography (SEC) (Chiba et al., 2022). In SEC, wild type KIF5A predominantly eluted at a single peak (Fig 3A blue shaded area). In addition, a small amount of wild type KIF5A was recovered from fractions that were eluted before the major peak and from the void volume (Fig 3A red and yellow shaded areas). SEC coupled with multi angle light scattering (SEC-MALS) analyses have shown that the main peak corresponds to KIF5A dimers and that the minor peak eluted before the main peak represents KIF5A oligomers (Chiba et al., 2022). Next, we examined KIF5A(Δexon27) and found that the Δexon27 mutation markedly changed the elution profile from that of wild type KIF5A. The major elution peak of Δexon27 shifted towards a higher molecular weight, which corresponds to oligomerization (Fig 3A red shaded area). Most KIF5A(Δexon27) was recovered from high molecular weight fractions but not from dimer fractions. Thus, the Δexon27 mutation induces oligomerization of KIF5A. We note that Δexon27 also increased the population recovered from the void volume, suggesting the potential of KIF5A(Δexon27) to form aggregates as well as oligomers (Fig 3A yellow shaded area).

**Figure 3.**
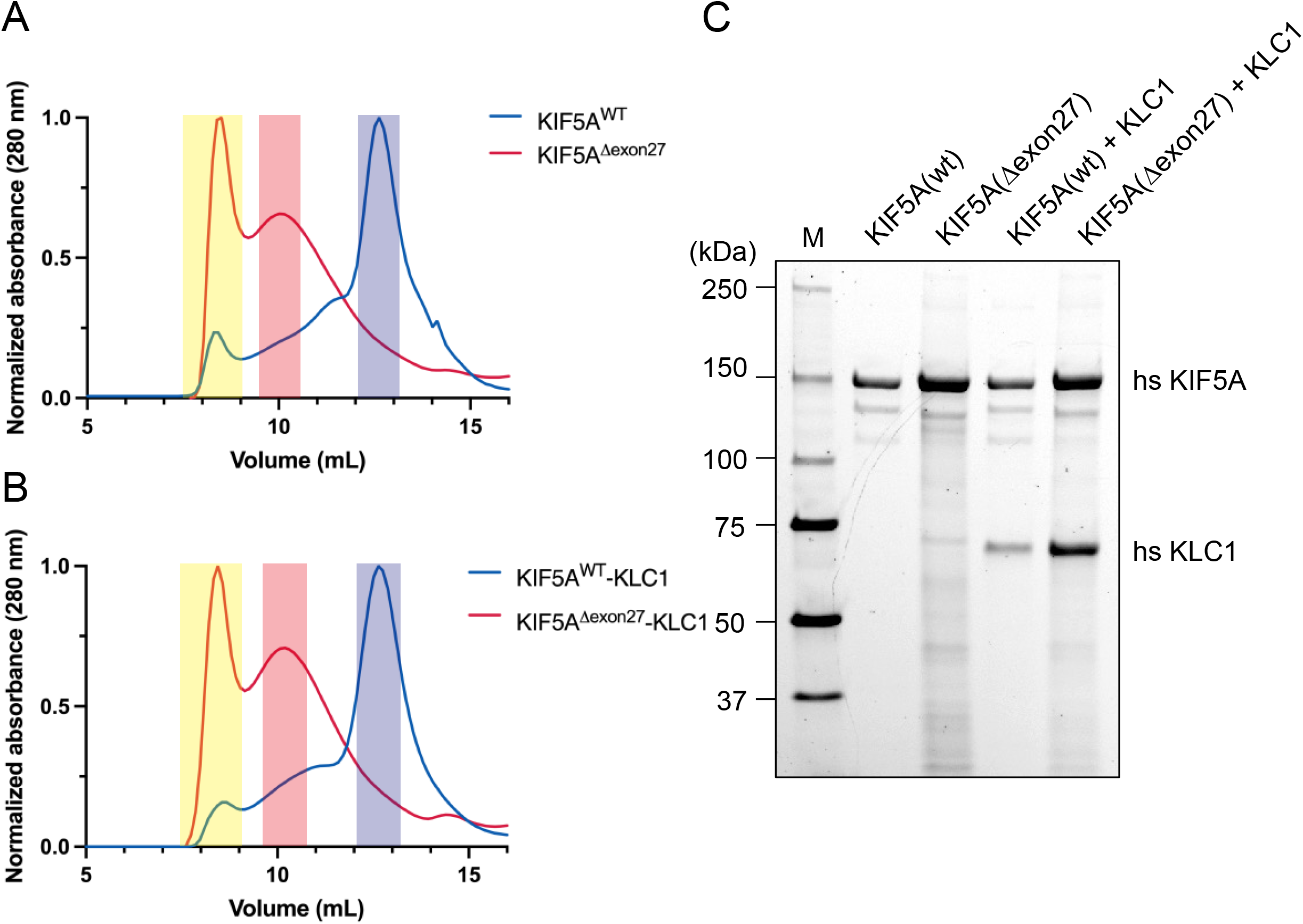
Purification of KIF5A and KIF5A(Δexon27) mScarlet was added to the C-terminus of either KIF5A (KIF5A::mSca) or KIF5A(Δexon27) (KIF5A(Δexon27)::mSca). Recombinant proteins were expressed in sf9 cells and then purified. (A and B) Normalized size exclusion chromatograms for KIF5A::mSca and KIF5A(Δexon27)::mSca (A) and KIF5A::mSca-KLC1 and KIF5A(Δexon27)::mSca-KLC1 (B). The yellow shaded area shows the void volume of the column corresponding to potential aggregates, the red shaded area shows the fraction corresponding to oligomers, and the blue shaded area shows the fraction corresponding to KIF5A dimers or KIF5A-KLC1 tetramers. (C) Coomassie blue stained gel showing purified KIF5A::mSca, KIF5A(Δexon27)::mSca, KIF5A::mSca-KLC1, and KIF5A(Δexon27)::mSca-KLC1. Wild type protein from the blue shaded fraction and Δexon27 protein from the red shaded fraction (A and B) were analyzed.

Next, to determine whether binding of the KLC subunit is affected, we co-expressed KLC1 with full length KIF5A or KIF5A(Δexon27). KLC1 was co-purified either with KIF5A(wt) or KIF5A(Δexon27) (Fig 3B and 3C). The ratio of heavy chains to light chains was almost 1:1 and was not markedly affected by Δexon27. Thus, binding to KLC is not abolished by the deletion of exon 27. The KIF5A-KLC1 complex showed an elution profile similar to that of KIF5A in the SEC analysis (Fig 3B blue lines). With KIF5A(Δexon27)-KLC1, we again observed a peak shifted towards a higher molecular weight and an increased population recovered from the void volume. These results suggest that the propensity of KIF5A(Δexon27) to form oligomers and aggregates is not largely affected by KLC1.

### KIF5A(Δexon27) oligomers are hyperactivated on microtubules

We analyzed the motility of purified microtubule motors at single-molecule resolution by total internal reflection fluorescence microscopy (Chiba et al., 2022; Chiba et al., 2019; McKenney, Huynh, Tanenbaum, Bhabha, & Vale, 2014). Purified full-length KIF5A(wt)::mSca and KIF5A(wt)::mSca-KLC1 did not bind or move well on microtubules (Fig 4A and 4B) because of the autoinhibitory mechanism described previously (Chiba et al., 2022; Coy, Hancock, Wagenbach, & Howard, 1999; Friedman & Vale, 1999; Hackney & Stock, 2000). In contrast, we found that full-length KIF5A(Δexon27)::mSca and full-length KIF5A(Δexon27)::mSca-KLC1 frequently bound to and moved along microtubules (Fig 4A–C). KLC1 suppresses the binding of KIF5A(Δexon27) onto microtubules (Fig 4C). Interestingly, Δexon27 mutation did not delete the IAK motif that is essential for autoinhibitory mechanism of kinesin heavy chain (Fig 1B)(Coy et al., 1999; Friedman & Vale, 1999; Hackney & Stock, 2000). The fluorescent intensities of KIF5A(Δexon27)::mSca and full-length KIF5A(Δexon27)::mSca-KLC1 were stronger than those of KIF5A(wt)::mSca and KIF5A(wt)::mSca-KLC1 (Fig 4B), suggesting the formation of oligomers. On average, the binding rates of KIF5A(Δexon27)::mSca and KIF5A(Δexon27)::mSca-KLC1 were nine times higher than those of KIF5A(wt)::mSca and KIF5A(wt)::mSca-KLC1, respectively (Fig 4C). We previously showed that the average run length and velocity of wild type KIF5A are 1.0 μm and 1.1 μm/sec, respectively (Chiba et al., 2022); however, we could not collect a sufficient number of samples to measure the run length and velocity of wild type proteins under the present condition. The median run lengths of KIF5A(Δexon27)::mSca and KIF5A(Δexon27)::mSca-KLC1 were approximately 3 μm each (Fig 4D), which is much longer than that of wild type KIF5A. No significant difference was detected between KIF5A(Δexon27)::mSca and KIF5A(Δexon27)::mSca-KLC1. The velocity of KIF5A(Δexon27)::mSca and KIF5A(Δexon27)::mSca-KLC1 was 0.4 μm/sec (Fig 4E and F), which is slower than that of wild type KIF5A. Taken together, these results suggest that KIF5A oligomers induced by Δexon27 are more active than those of wild type KIF5A both in the presence and absence of the KLC subunit (Table 1).

**Table 1.**
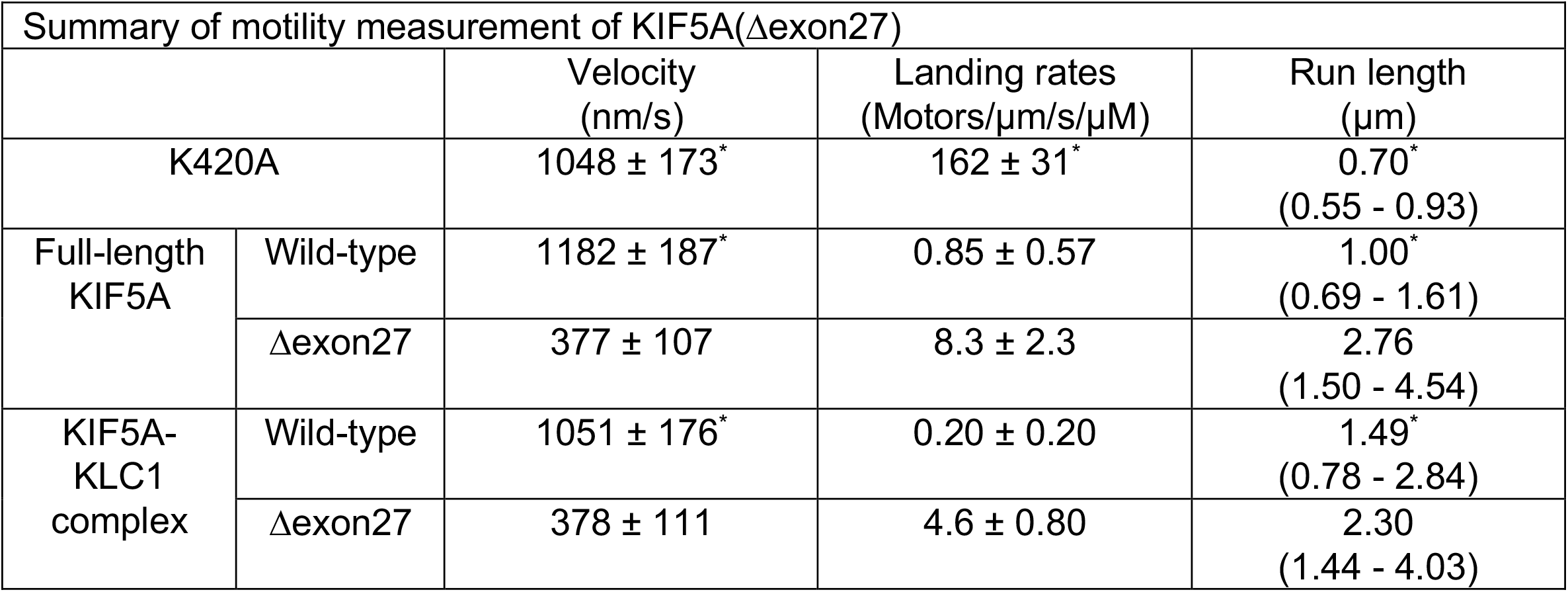
Measured Parameters of Human Kinesin Isotypes. The motility parameters of tail-truncated K420s, full-length KIF5A dimers and KIF5-KLC1 tetramers. Velocity represents mean ± SD calculated from Gaussian fitting based on bootstrap distribution. Landing rate represents total processive and non-processive microtubule binding events over microtubule length, time and motor concentration in the presence of ATP. Run length represents median and interquartile range. Processivity represent frequency (%) of molecules showing directional movement over the total microtubule binding events. ^*^ Values are described in Chiba et al (2022).

**Figure 4.**
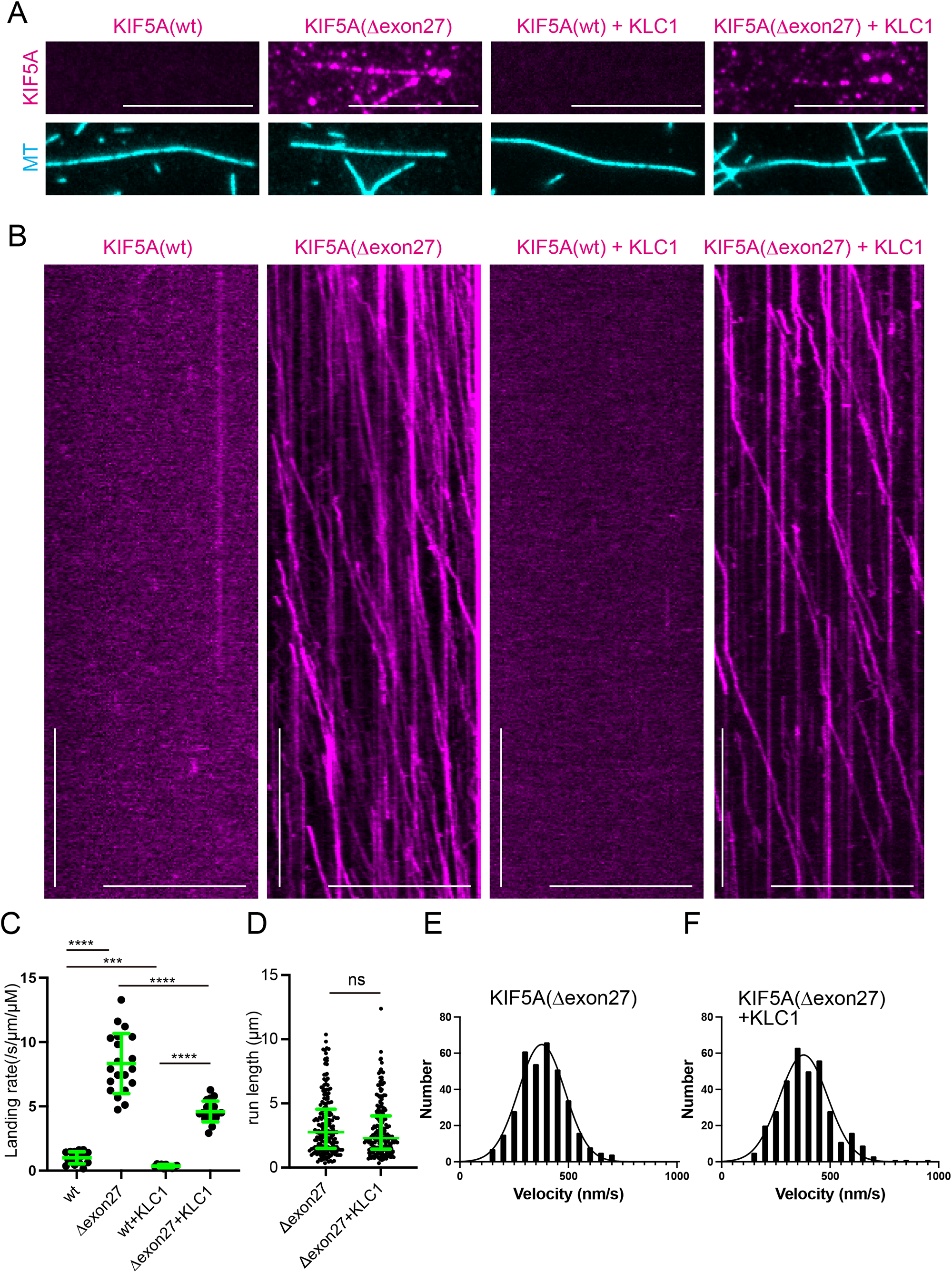
Motility of amyotrophic lateral sclerosis-associated KIF5A on microtubules. The motility of 10 nM KIF5A::mSca, 10 nM KIF5A(Δexon27)::mSca, 10 nM KIF5A::mSca-KLC1 complex, and 10 nM KIF5A(Δexon27)::mSca-KLC1 complex on microtubules in the presence of 2 mM ATP was observed by TIRF microscopy. Wild type protein from the blue shaded fraction and Δexon27 protein from the red shaded fraction in Figure 3A and B were observed. (A) Representative TIRF images of KIF5A (magenta) and microtubules (cyan). Note that KIF5A(Δexon27)::mSca and KIF5A(Δexon27)::mSca-KLC1 bound to microtubules more frequently than KIF5A::mSca and KIF5A::mSca-KLC1. Bars, 10 μm. (B) Representative kymographs of KIF5A::mSca, KIF5A(Δexon27)::mSca, KIF5A::mSca-KLC1 complex, and KIF5A(Δexon27)::mSca-KLC1 complex. Scale bars represent 10 sec (vertical) and 10 μm (horizontal). (C) Dot plots showing the landing rates of KIF5A::mSca (wt), KIF5A(Δexon27)::mSca (Δexon27), KIF5A::mSca-KLC1 complex (wt+KLC1), and KIF5A(Δexon27)::mSca-KLC1 complex (Δexon27+KLC1). The number of KIF5A oligomers that bound to microtubules was counted and adjusted by the time window, microtubule length, and concentration. Each dot shows one data point. N = 20 kymographs from each sample. Data were analyzed by Brown-Forsythe and Welch ANOVA followed by Dunnett’s multiple comparisons test. ***, adjusted P value < 0.001. ****, adjusted P value < 0.0001. (D) Dot plots showing the run length of KIF5A(Δexon27)::mSca (Δexon27) and KIF5A(Δexon27)::mSca-KLC1 complex (Δexon27+KLC1). N = 202 (Δexon27) and 201 (Δexon27+KLC1). Data were analyzed with the Mann-Whitney test. ns, adjusted P value > 0.05 and not statistically significant (E and F) Gaussian fit and histogram showing the velocity of KIF5A(Δexon27) and the KIF5A(Δexon27)::mSca-KLC1 complex (377 ± 107 nm/sec (KIF5A(Δexon27)) and 378 ± 111 nm/sec (KIF5A(Δexon27)-KLC1)). The means ± standard deviation were obtained from best-fit values. N = 349 and 337, respectively. The velocity of KIF5A(Δexon27) was not affected by KLC1. Note that a sufficient number of samples could not be collected for KIF5A(wt)::mSca and KIF5A(wt)::mSca-KLC1.

### KIF5A(Δexon27) oligomers tend to form aggregates

To examine the properties of KIF5A(Δexon27), we incubated purified mScarlet, KIF5A::mSca and KIF5A(Δexon27)::mSca at 37°C and measured the turbidity of protein solutions by determining the optical density at 600 nm, which was used to monitor the formation of protein aggregates in solution in a previous study (Schafheimer & King, 2013). Immediately before incubation, protein solutions were centrifuged and clarified. The turbidity of purified KIF5A(Δexon27)::mSca increased gradually while those of mScarlet and KIF5A::mSca did not (Fig 5A). Twenty-four hours later, the solution was again centrifuged, and purified mSca::KIF5A(Δexon27) formed a protein pellet (Fig 5B). At this concentration, purified mScarlet and KIF5A::mSca did not form visible pellets at all, even after the 24-hour incubation. These data indicate that purified KIF5A(Δexon27) is predisposed to form protein aggregates *in vitro*.

**Figure 5.**
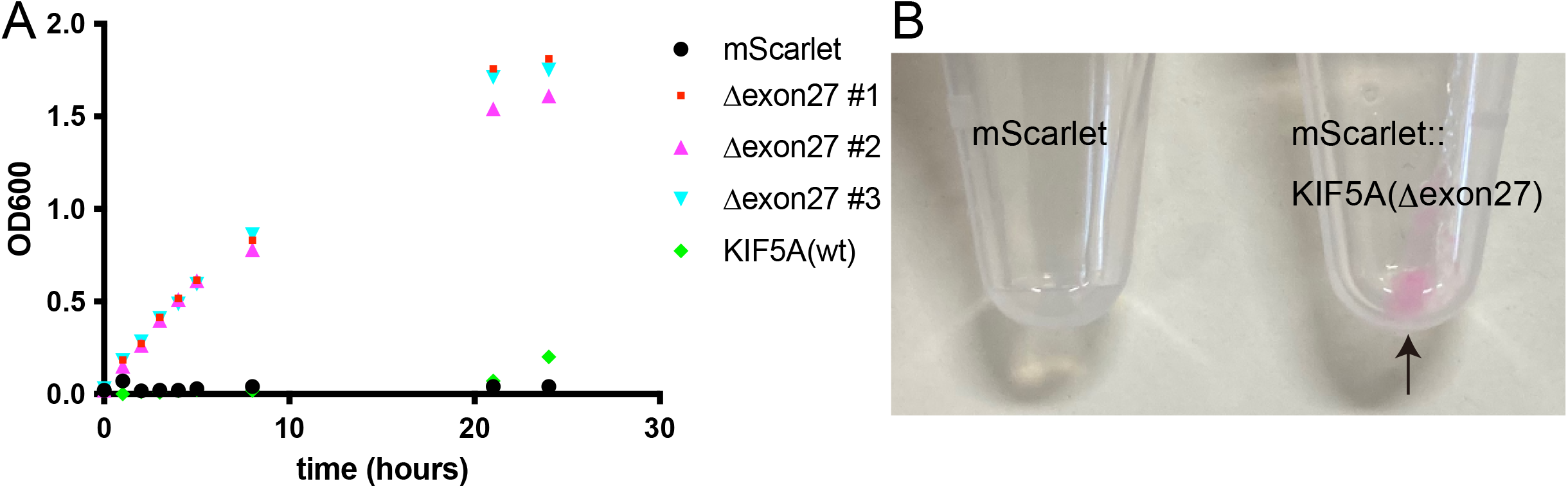
Aggregate formation *in vitro*. Purified mScarlet, KIF5A::mSca or KIF5A(Δexon27)::mSca (1 μM) was incubated at 37°C. (A) Turbidity of the solution. The optical density of samples was measured at a wavelength of 600 nm (OD600) and plotted at the indicated time points. Results from three independent experiments are shown for KIF5A(Δexon27)::mSca.. (B) After 24 hours of incubation, the samples were centrifuged at 18,000 g for 10 min, and the presence of a pellet was detected. Note that a protein pellet was induced in the KIF5A(Δexon27)::mSca sample, while no pellet was observed in purified mScarlet.

### KIF5A(Δexon27) is toxic in C. elegans neurons

Previous studies have shown that ALS is caused by toxic gain-of-function mutations (Bruijn et al., 1998; Johnson et al., 2009; Kwiatkowski et al., 2009). Therefore, we examined the toxicity of KIF5A(Δexon27) in *C. elegans* neurons. The morphology of mechanosensory neurons was compared after expressing human KIF5A and KIF5A(Δexon27) (Fig 6A–F). Mechanosensory neurons were analyzed because many studies have shown that these neurons have very stereotypical morphology and show small variations in the wild type background (Gallegos & Bargmann, 2004; Ghosh-Roy, Goncharov, Jin, & Chisholm, 2012).The morphology of posterior lateral microtubule (PLM) and posterior ventral microtubule (PVM) neurons was observed and compared in day 1 adults. PLM neurons had a long straight axon along the body in wild type cells (Fig 6A and B). The PVM neuron extends an axon that grows ventrally and then turns anteriorly when it reaches the ventral nerve cord (Fig 6A and B). No significant differences were found between KIF5A-expressing neurons and control neurons (Fig 6C). In contrast, KIF5A(Δexon27)-expressing neurons showed morphological defects (Fig 6D–F). Approximately 60% of worms showed abnormal PLM and PVM morphologies (Fig 6D), but no stereotypical abnormalities were found. The PLM cell body was often mislocalized, and some worms had PLM neurons with multiple neuronal processes, while other worms had PLM neurons with bent axons. Approximately 20% of worms showed neuronal loss (Fig 6E). These data suggest that KIF5A(Δexon27) is toxic in neurons.

**Figure 6.**
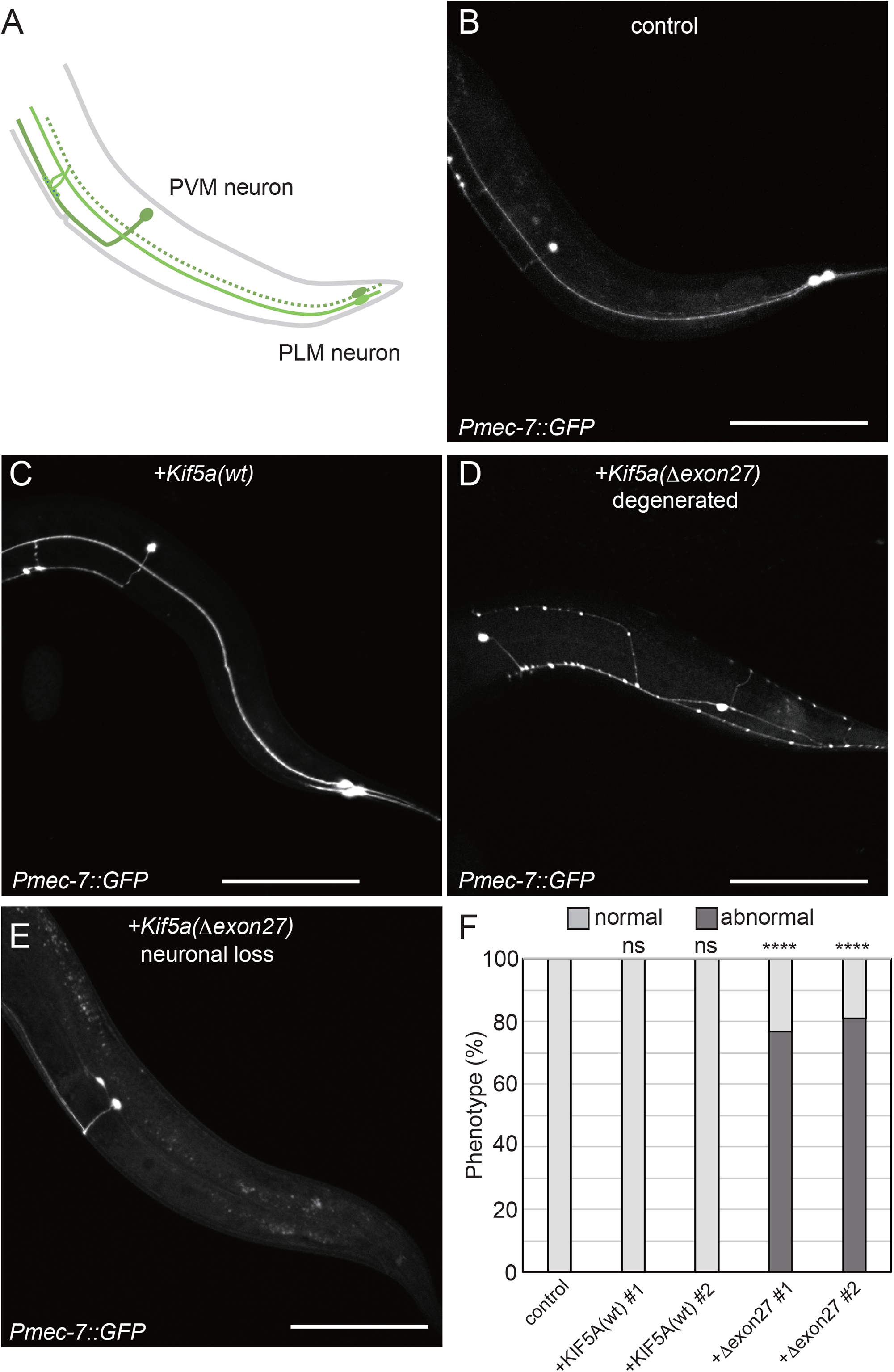
Neuronal toxicity in *Caenorhabditis elegans*. Human KIF5A or KIF5A(Δexon27) was expressed in a *C. elegans* strain expressing green fluorescent protein (GFP) in mechanosensory neurons. (A and B) Morphology of posterior ventral microtubule (PVM) and posterior lateral microtubule (PLM) mechanosensory neurons. A schematic drawing of PVM and PLM neurons (A) and a representative image (B) of PVM and PLM neurons visualized by GFP expressed under the *mec-7* promoter (*Pmec-7*) are shown. (C–E) Representative images showing *Kif5a*-expressing mechanosensory neurons (C) and *Kif5a(*Δ*exon27)*-expressing neurons (D and E). (D) The formation of beads along the axon, an injured axon marker, is seen. Moreover, the cell body of PLM neuron is anteriorly mislocalized. The axon of PLM neuron abnormally bends while the axon of PVM turns posteriorly. (E) PLM neuron is lost. Bars, 100 μm. (F) *C. elegans* phenotypes were classified according to mechanosensory neuron morphology into two categories: normal morphology (light gray) and abnormal morphology (dark gray), and the number of worms in each category was counted. N = 87 (control strain), 68 (KIF5A(wt)-expressing strain #1), 75 (KIF5A(wt)-expressing strain #2), 73 (KIF5A(Δexon27)-expressing strain #1), and 79 (KIF5A(Δexon27)-expressing strain #2). Data were analyzed with a chi-square test with Bonferroni correction. ns, adjusted P value > 0.05 and not statistically significant, ****, adjusted P value < 0.0001; compared with the control strain.

## Discussion

Previous studies have suggested that KIF5A(Δexon27) causes ALS by a loss-of-function mechanism because the cargo-binding tail domain is deleted (D. Brenner et al., 2018; Nicolas et al., 2018). However, while several loss-of-function mutations have been found in the motor domain of KIF5A, these mutations cause SPG, rather than ALS (Ebbing et al., 2008; Reid et al., 2002). None of the motor domain mutations in KIF5A have been associated with ALS. These genetic data suggest that ALS mutations in KIF5A are not simple loss-of-function mutations. Protein aggregates are often associated with neurodegenerative disorders (Soto, 2003), and numerous ALS-associated mutations have been identified in other genes such as TARDBP (TDP-43 gene), SOD, and FUS (Aoki et al., 1993; Kabashi et al., 2008; Kwiatkowski et al., 2009; Vance et al., 2009). These mutations commonly induce aggregates that are toxic in cells (Bruijn et al., 1998; Johnson et al., 2009; Kwiatkowski et al., 2009). Our data suggest that the Δexon27 mutation in KIF5A induces toxic aggregates. Furthermore, while we were preparing this manuscript, a preprint supporting the same conclusion was posted on Biorxiv (Pant et al., 2022). The study showed that KIF5A(Δexon27) causes aggregate formation in cultured cells and is toxic when expressed in *Drosophila*. The study also showed that unpurified KIF5A(Δexon27) in cell lysates is more active than wild type KIF5A. What induces the formation of KIF5A(Δexon27) aggregates? It is possible that unidentified proteins bind to the abnormal C-terminus of KIF5A(Δexon27) and induce hyperactivation and aggregation. However, our assays using purified proteins strongly suggest that KIF5A(Δexon27) is hyperactive and is predisposed to form aggregates without the involvement of other factors.

The Δexon27 mutation induces hyperactivation of KIF5A. KIF5 is inhibited by KLC-dependent and independent autoinhibitory mechanisms (Chiba et al., 2022; Hackney & Stock, 2000; Verhey et al., 1998). Binding with cargo vesicles or cargo complex unlock the autoinhibition. The binding of KIF5A(Δexon27) onto microtubules is suppressed by KLC1 (Fig 4C), suggesting that KLC-dependent autoinhibitory mechanism works even in KIF5A(Δexon27). Interestingly, the IAK motif is not affected by the Δexon27 mutation (Fig 1B). It was shown that hydrophobic materials such as polystyrene beads and glass surface can mimic cargos and activate KIF5-KLC complex when they bind to the tail region (Vale et al., 1985). Δexon27 mutation induces the formation of large oligomers (Fig 3A and B). The large KIF5A(Δexon27) oligomer may mimic cargos and activate the motor. It is also possible that Δexon27 mutation disrupts previously unknown autoinhibitory mechanisms.

What induces toxicity in neurons? Toxicity in worm neurons would help to clarify neurotoxic mechanisms. One possibility is that hyperactivated KIF5A changes the distribution of cargo organelles and induces cellular toxicity. We have shown that another kinesin, human KIF1A, is functional in worm neurons (Chiba et al., 2019). We show here KIF5(Δexon27) binds to the cargo-binding adaptor KLC1 (Fig 3C). Thus, cargo transport may be affected by KIF5(Δexon27) even in worm neurons. Another possibility is that cargo transport is not related and KIF5(Δexon27) aggregates are toxic in neurons as is the case in other ALS-associated mutations (Bruijn et al., 1998; Johnson et al., 2009; Kwiatkowski et al., 2009). Aggregates change cellular metabolisms and induces neuronal death (Soto, 2003). These possibilities will be discriminated by analyzing cargo transport and distributions in worm neurons. If aggregates are toxic, hyperactivation of KIF5A motor may enhance the toxicity because hyperactivation causes mis-accumulation of KIF5A at axonal tips, leading to a high concentration of KIF5A and aggregate formation. This hypothesis is supported by the observation that KIF5(Δexon27) aggregates were often found at neurite tips in CAD cells (Fig 1).

We have previously shown that KIF5A forms more oligomers than KIF5B and KIF5C *in vitro* (Chiba et al., 2022). Additionally, KIF5A oligomers are more active than KIF5A dimers. However, the physiological significance of these phenomena remains elusive. We show here that even wild type KIF5A has a propensity to form more aggregates in the cell than KIF5B (Fig 1). A similar property has been found for TDP-43. TDP-43 is intrinsically aggregation-prone, which implies that it may be directly involved in the pathogenesis of sporadic ALS (Johnson et al., 2009). The propensity of KIF5A to oligomerize may be related to sporadic ALS and may be enhanced by ALS-associated KIF5A mutations.

## Experimental Procedures

### Molecular biology

Polymerase chain reaction (PCR) was performed using KOD FX neo (TOYOBO, Tokyo, Japan) as described in the manual. Restriction enzymes were purchased from New England BioLabs Japan (Tokyo, Japan).

pAcebac1 plasmids containing human *KIF5A*(BC146670) human *KIF5B*(BC126281), and *KLC1* (BC008881) was described previously (Chiba et al., 2022).

To generate mScarlet::KIF5A and mScarlet::KIF5B expressing plasmids, pmScarletC1 plasmid was obtained from Addgene. KIF5A and KIF5B were amplified by PCR and transferred to pmScarletC1. To generate an EGFP::KIF5B expressing vector, EGFP was amplified from pEGFPN1::KIF1A (Niwa et al., 2008) and replaced with mScarlet by using AgeI and XhoI sites. Δexon27 mutation was introduced by Gibson assembly. cDNA fragment encoding the mutated domain (Fig 1A) was synthesized by gBlocks (Integrated DNA Technologies Japan, Tokyo, Japan) and replated with the wild-type fragment by Gibson assembly (Gibson et al., 2009). Plasmids used in this paper is described in ***supplementary table S1***. Key plasmids were deposited to Addgene.

### CAD cell experiments

CAD cells were obtained from European Collection of Cell Cultures (Salisbury, UK) and maintained as described (Qi et al., 1997). For observation, cells were cultured on glass coverslips (Matsunami, Tokyo, Japan). Plasmid transfection was performed by Lipofectamine LTX (Thermo Fisher Scientific) as described in the manufacturer’s manual. 24 hours after the transfection, mScarlet and GFP signals were observed under Zeiss Axio Observer inverted microscope equipped with LSM800 confocal unit (Carl Zeiss). x40 water immersion objective lens (Numerical Aperture: 1.1) was used for imaging. ZEN software (Carl Zeiss) was used to control the system.

### Purification of KIF5A

pAcebac plasmids were transformed to generate bacmid. Sf9 cells were maintained as a suspension culture in Sf-900II serum-free medium (SFM) (Thermo Fisher Scientific) at 27°C. To prepare baculovirus, 1 × 10^6^ cells of Sf9 cells were transferred to each well of a tissue-culture treated 6 well plate. After the cells attached to the bottom of the dishes, about ∼5 μg of bacmid were transfected using 6 μl of Cellfectin II reagent (Thermo Fisher Scientific). 5 days after initial transfection, the culture media were collected and spun at 3000 x g for 3 min to obtain the supernatant (P1). For protein expression, 400 ml of Sf9 cells (2 × 10^6^ cells/ml) were infected with 100 μl of P1 virus and cultured for 65 hr at 27°C. Cells were harvested and resuspended in 25 ml of lysis buffer (50 mM HEPES-KOH, pH 7.5, 150 mM KCH_3_COO, 2 mM MgSO4, 1 mM EGTA, 10% glycerol) along with 1 mM DTT, 1 mM PMSF, 0.1 mM ATP and 0.5% TritonX-100. After incubating on ice for 10 min, the lysates were centrifuged at 15000 x g for 20 min at 4°C. The resulting supernatant were subject to affinity chromatography described below.

For affinity chromatography, the supernatants were put over a column of Streptactin XT resin (IBA) at 4°C. The columns were then washed with excess lysis buffer to remove unbound material and the proteins were eluted in lysis buffer containing 100 mM D-biotin. Eluted proteins were further purified via size exclusion chromatography using a Superose 6 10/300 GL column (Cytiva) equilibrated in lysis buffer. Fractions containing the proteins were combined and concentrated on amicon spin filters with a 50 kDa cutoff. Concentrated proteins were frozen in LiN_2_ and stored at -80°C.

### TIRF single-molecule motility assays

TIRF assays were performed as described (Chiba et al., 2019). Tubulin was purified from porcine brain as described (Castoldi & Popov, 2003). Tubulin was labeled with Biotin-PEG_2_-NHS ester (Tokyo Chemical Industry, Tokyo, Japan) and AZDye647 NHS ester (Fluoroprobes, Scottsdale, AZ, USA) as described (Al-Bassam, 2014). To polymerize Taxol-stabilized microtubules labeled with biotin and AZDye647, 30 μM unlabeled tubulin, 1.5 μM biotin-labeled tubulin and 1.5 μM AZDye647-labeled tubulin were mixed in BRB80 buffer supplemented with 1 mM GTP and incubated for 15 min at 37°C. Then, an equal amount of BRB80 supplemented with 40 μM taxol was added and further incubated for more than 15 min. The solution was loaded on BRB80 supplemented with 300 mM sucrose and 20 μM taxol and ultracentrifuged at 100,000 g for 5 min at 30°C. The pellet was resuspended in BRB80 supplemented with 20 μM taxol. Glass chambers were prepared by acid washing as previously described (McKenney et al., 2014). Glass chambers were coated with PLL-PEG-biotin (SuSoS, Dübendorf, Switzerland). Polymerized microtubules were flowed into streptavidin adsorbed flow chambers and allowed to adhere for 5–10 min. Unbound microtubules were washed away using assay buffer [90 mM Hepes pH 7.4, 50 mM KCH_3_COO, 2 mM Mg(CH_3_COO)_2_, 1 mM EGTA, 10% glycerol, 0.1 mg/ml biotin–BSA, 0.2 mg/ml kappa-casein, 0.5% Pluronic F127, 2 mM ATP, and an oxygen scavenging system composed of PCA/PCD/Trolox. Purified motor protein was diluted to indicated concentrations in the assay buffer. Then, the solution was flowed into the glass chamber. An ECLIPSE Ti2-E microscope equipped with a CFI Apochromat TIRF 100XC Oil objective lens, an Andor iXion life 897 camera and a Ti2-LAPP illumination system (Nikon, Tokyo, Japan) was used to observe single molecule motility. NIS-Elements AR software ver. 5.2 (Nikon) was used to control the system.

### Worm experiments and strains

*C. elegans* strains were maintained as described previously (S. Brenner, 1974). Wild-type worm N2 and feeder bacteria OP50 were obtained from C. elegans genetic center (CGC) (Minneapolis, MN, USA). Nematode Growth Media (NGM) agar plates were prepared as described (S. Brenner, 1974). Transformation of C. elegans was performed by DNA injection as described (Mello, Kramer, Stinchcomb, & Ambros, 1991).

Strains used in this study are described in supplementary table S2.

### Analysis of mechanosensory neurons in C. elegans

Mechanosensory neurons were visualized using *uIs31* [*Pmec-7::gfp*] marker that was obtained from *C. elegans* genetic center (Minnesota, USA) (Chalfie, Tu, Euskirchen, Ward, & Prasher, 1994). Strains expressing human KIF5A or KIF5A(Δexon27) were observed under Zeiss Axio Observer inverted microscope equipped with LSM800 confocal unit (Carl Zeiss, Jena, Germany). x20 objective lens (Numerical Aperture: 0.8) was used for imaging. ZEN software (Carl Zeiss) was used to control the system. Fiji was used to analyze image files (Schindelin et al., 2012).

## Acknowledgements

We thank all of the members of the McKenney lab (UC Davis), Sugimoto lab (Tohoku Univ.), and Niwa lab (Tohoku Univ.) for helpful discussions. SN was supported by JSPS KAKENHI (20H03247, 19H04738, 20K21378), the Kato Memorial Bioscience Foundation, and Takeda Science Foundation. KC was supported by JSPS KAKENHI (21K20621) and MEXT Leading Initiative for Excellent Researchers. Some *C. elegans* strains and OP50 were obtained from the CGC. We thank Lisa Kreiner, PhD, from Edanz (https://www.jp.edanz.com/ac) for editing a draft of this manuscript.

