## Supplemental Table S1 for "An ALS-associated KIF5A mutant forms oligomers and aggregates and induces neuronal toxicity"

### Supplementary Table S1 Plasmid list

| Plasmid | Description | Backbone | Addgene ID |
|---|---|---|---|
| pEGFPC1 | pEGFPC1 obtained from Clontech | | |
| pmScarletC1 | pmScarletC1 obtained from Addgene | | |
| pSN837 | CMV promoter::mScarlet::human KIF5A | pmScarletC1 | 184825 |
| pSN838 | CMV promoter::mScarlet::human KIF5A(Δexon27) | pmScarletC1 | 184826 |
| pSN805 | CMV promoter::EGFP::human KIF5A. mScarlet in pSN837 was replaced with EGFP. | pSN837 | |
| pSN738 | CMV promoter::EGFP::human KIF5B. | pEGFPC1 | |
| pSN816 | *mec-7* promoter::human KIF5A | pSM vector obtained from Dr. Kang Shen | |
| pSN817 | *mec-7* promoter::human KIF5A(Δexon27) | pSN816 | |
| pMOM657 | pAcebac1 human KIF5A::mScarlet::StrepII. Described in Chiba et al. (2022). | pAcebac1 | |
| pSN526 | pAcebac1 human KIF5A(Δexon27)::mScarlet::2xStrepII | pAcebac1 | 184827 |
| pMOM489 | pIDS His tag::FLAG::human KLC1. Described in Chiba et al. (2022). | pIDS | |
| pMOM659 | Recombined plasmid. pAcebac1 human KIF5A::mScarlet::StrepII + pIDS His::FLAG::human KLC1. Described in Chiba et al. (2022). | pAcebac1 and pIDS | |
| pSN840 | Recombined plasmid. pAcebac1 human KIF5A(Δexon27)::mScarlet::2xStrepII + pIDS His::FLAG::human KLC1 | pAcebac1 and pIDS | 184832 |
