## Supplemental Table S2 for "An ALS-associated KIF5A mutant forms oligomers and aggregates and induces neuronal toxicity"

Supplementary Table S2 Strain list

| ***strain*** | ***genotype*** | ***description*** |
|---|---|---|
| *TU2769* | *uIs31[Pmec-7::gfp] III* | |
| *OTL155* | *uIs31; jpnEx110[Pmec-7::KIF5A]* | pSN816 was injected to TU2769. |
| *OTL156* | *uIs31; jpnEx111[Pmec-7::KIF5A]* | pSN816 was injected to TU2769. |
| *OTL157* | *uIs31; jpnEx112[Pmec-7::KIF5A(Dexon27)]* | pSN817 was injected to TU2769. |
| *OTL158* | *uIs31; jpnEx113[Pmec-7::KIF5A(Dexon27)]* | pSN817 was injected to TU2769. |
